# Cortical Spectral Dynamics in Flow and Frustration

**DOI:** 10.64898/2026.09.21.753264

**Authors:** Raymond P. Viviano, Shailesh N. Joshi, Ercan M. Dede

## Abstract

Flow is a subjective state where tasks feel effortless and rewarding. It has been linked to distinct neural signatures, yet dynamics of these signals remain poorly understood. Here, we investigated spectral dynamics of brain activity in flow. We recorded electroencephalogram (EEG) data at positions AF7, AF8, TP9, and TP10 using a lightweight wearable device worn by ten adult participants while they completed a go-signal task and two stop-signal tasks. Task difficulty was continuously adjusted to induce either a sense of flow or frustration. Time-resolved spectral power and interregional coherence were extracted with continuous wavelet and wavelet coherence transforms; subjective flow was assessed after each block with the short-form Flow-State Scale. Subjective ratings confirmed that the flow and frustration conditions diverged in perception of flow. In the flow block, errors on ‘stop’ trials coincided with pre-stimulus reductions in right temporoparietal gamma power and frontal beta and gamma power, alongside diminished frontotemporal gamma coherence. Comparison of flow and frustration revealed higher temporal beta power during frustration, consistent with greater cognitive load. Flow may involve finely balanced beta and gamma dynamics and brief collapses of these rhythms may predict task errors. Spectral dynamics are actionable features for closed-loop engagement monitoring applications.

## 1. Introduction

**F**LOW is a psychoaffective state of effortless concentration where an individual is absorbed in a task while actions occur smoothly [1], [2]. It has positive valence, with individuals experiencing intrinsic reward. Generally, flow arises from activities when there is a balance between challenge and skill, such as playing a musical instrument [3], a sport [4], or a video game [5], [6]. However, flow experiences have also been reported for tasks in lab settings, including mental arithmetic [7]–[9], which may not be a pleasurable pastime for most. While flow experiences have been well characterized in psychological literature, neuroscientific investigation remains sparse [10]. Regardless, competing hypotheses have emerged, including Transient Hypofrontality, Synchronization, and Neural Proficiency hypotheses.

### A. The Transient Hypofrontality Hypothesis

The Transient Hypofrontality hypothesis [11] posits that flow arises from decreased processing in an explicit, verbalizable knowledge system to free up resources for implicit, skills-based processing. While in a flow-state, per this hypothesis, analytical self-monitoring by the frontal lobes diminishes to enable automatic subcortical task execution (e.g., via the basal ganglia) [11]. Some evaluations have supported a hypofrontality account. An fNIRS study found that higher self-reported flow correlated with lower prefrontal oxygenated hemoglobin during a verbal fluency task [12]. Furthermore, perfusion MRI has revealed decreased medial prefrontal (mPFC) blood flow during a flow experience compared to boredom and overload [8]. Decreased Default Mode Network (DMN) activity, including mPFC, posterior cingulate cortex, and medial temporal regions, has also been reported in flow compared to boredom and overload with fMRI [9]. However, while mPFC activity was lower during flow, task-positive dorsolateral prefrontal cortex (dlPFC) activity was greater at the same time, failing to support a strict hypofrontality interpretation. Furthermore, DMN activity suppression alongside Central Executive Network activity increases is a common occurrence during external task demands [13], [14] and are not sufficient observations to support the Transient Hypofrontality Hypothesis.

Despite some support for reduced frontal activity [8], [12], other findings show that prefrontal processes are not subdued in flow. In addition to [9], some studies observed sustained or enhanced frontal activity. For example, a study of flow with a mental arithmetic task found that frontal midline EEG theta power (an index of focused attention and executive control) was just as high during flow as during overload, and higher than in boredom [7]. Broader evidence from altered flow-like states such as meditation also show strong prefrontal activity in neuroimaging, not suppression [15], [16]. The hypofrontality hypothesis also separates the experience of flow from creativity, which recruits prefrontal circuits [17] and involves critical evaluation of novelty. Thus, the hypothesis may overemphasize automaticity, favoring overlearned motor skills over cognitive tasks, and may not easily explain flow experiences involving discovery and novelty.

### B. The Synchronization Hypothesis

The Synchronization Hypothesis was originally formulated in the context of media use and posits that when skill and task challenge are matched, brain networks for attention and reward processing synchronize [18]–[20]. Flow may emerge from functional connectivity between systems that focus attention (e.g., frontal and parietal control regions) and those that signal intrinsic reward (e.g., striatal dopamine circuits), producing a distinct brain state perceived as effortless and enjoyable.

An fMRI study explicitly tested synchronization predictions; participants played a video game with low, high, or optimally-balanced challenge. Functional connectivity between cognitive control (dlPFC) and reward (ventral striatal) regions was higher during the flow condition compared to boredom and overload [20]. In addition, self-report confirmed that the balanced condition elicited high intrinsic reward and engagement. By contrast, when difficulty did not match skill, connectivity between these networks dropped and DMN regions became more active, signaling disengagement. In other fMRI research, flow induced by playing video games was reported to be accompanied by co-activation of frontoparietal attention and dopaminergic midbrain regions [21], reinforcing the idea of simultaneous engagement of cognitive control and reward circuitry in flow. Other fMRI research has found flow to increase activity in the right inferior frontal gyrus, anterior insula, and midbrain [9]. The anterior insula and inferior frontal gyrus are involved in attention and task monitoring [22], [23], while midbrain and basal ganglia are central to reward and motivation [24], [25]. While not functional connectivity analyses, joint activations of regions during flow suggests that task and reward networks may work in concert.

Direct evidence for network synchronization in flow is still emerging; studies explicitly examining functional connectivity are limited. Regardless, many researchers have proposed unified models of synchronization and hypofrontality [10], [15], [26]; the flow experience may involve a complex pattern of frontal activity increases and decreases that reduce self-reflection while maintaining executive function. Concurrently, the brain may achieve flow by silencing task-disruptive parts of the prefrontal cortex (PFC) while synchronizing control circuits with reward pathways to sustain focus and enjoyment.

### C. The Neural Proficiency Hypothesis

While more in the ‘expert performance’ rather than ‘flow-state’ literature, neural proficiency reflects flexible adaptation of brain activity between efficient automatic processing and effortful controlled processing, in expert performers, to meet situational demands [27], [28]. Neural Proficiency suggests that optimal performance involves energetically efficient neural processing that requires prior skill proficiency. It emphasizes the capability to maintain high performance through regulation of higher-order cognitive and attentional resources, not just minimal energy expenditure. When an individual is skilled, the brain may execute a task with some degree of automaticity, leading to a sense of effortlessness. Experts often show lower overall brain activity alongside elevated activity in task-relevant regions compared to novices [29], [30].

Certain brain regions, e.g., basal ganglia and cerebellum, are involved in automated, overlearned actions that can be overridden by goal-directed actions determined by the PFC [31], [32]. Skill mastery may enable task allocation to these efficient neural circuits, requiring minor oversight from central-executive regions [26]. Thus, task execution may be somewhat automatic and energetically efficient but still require cognitive effort to meet overarching ambitions. Anterior regions are involved but only at a necessary minimum level, similar to the transient hypofrontality hypothesis. Neural proficiency is supported by some research on athletic practice. Evaluation of golfers found that increased temporal alpha power and reduced alpha connectivity between temporal and frontal areas mediated putting performance improvements [33]. Alpha oscillations may help structure cortical operation through the targeted suppression of regional activity; i.e., alpha may guide the distribution of resources, shifting activity away from areas with higher alpha power toward regions with lower alpha power [34], [35]. However, there is room to debate if skill proficiency is necessary to experience flow. As flow is defined in part by a balance of challenge and skill, novices and experts may both experience it as long as challenges are appropriate for specific skill levels. As with the transient hypofrontality hypothesis, a neural proficiency hypothesis may overemphasize the “automaticity of action” aspect of flow.

### D. Prior EEG Spectral Findings During Flow

Across mental-arithmetic and writing paradigms, the electrophysiological signature of flow involves a rise in mid-frontal theta and an inverted-U modulation of mid-frontal and temporal alpha. Katahira et al. found that flow was characterized by elevated frontocentral theta together with moderate alpha that tracked perceived task balance and concentration [7]. Knierim et al. replicated this, showing that temporal alpha power follows a quadratic trajectory where low or high values reflect sub-optimal engagement [36], [37]. These findings suggest that theta activity reflects sustained executive maintenance [38] in flow, whereas moderate alpha reflects an optimal coupling of attention resources and working-memory load that supports a subjective sense of effortlessness.

Dual-task designs elucidate how flow directs processing away from irrelevant input. During video game induced flow, suppression of the P3 response in parietal cortex to oddball tones occurred alongside increased mid-frontal alpha, and the magnitude of this alpha rise predicted quick response times [5]. In another gaming paradigm, trials self-rated as high in flow elicited attenuated auditory evoked potentials and heightened beta oscillations in the ACC and temporal pole; top-down beta connectivity from ACC to primary motor cortex also scaled positively with subjective absorption [6]. Flow may involve top-down alpha-mediated inhibition of extraneous stimuli coupled with beta-driven reinforcement of frontocortical control, facilitating goal-directed behavior while minimizing distraction. In general, the EEG flow-state research suggests frontal-midline theta engagement, context-sensitive alpha gating, and task-specific beta activity.

#### E. Hypotheses and Contribution

While prior analyses evaluated functional connectivity, task-related activity changes, and EEG spectra in flow, EEG dynamics still remain underexplored. Therefore, we evaluated dynamics in spectral power and coherence in frontal and temporoparietal leads between a go-signal task, and a stop-signal task with flow- and frustration-inducing conditions, using a lightweight Muse 2 EEG headset [39]. Overall, while the major hypotheses on the neural activity patterns that produce flow have some literature support, recent reviews highlight inconsistencies [10], [40]. Neuroscientific evaluation is still too sparse to formulate a unifying theory that addresses the aforementioned hypotheses; and some of the results in the extant literature do not fit neatly into these frameworks. Regardless, our hypotheses were informed by these ideas as well as specific EEG results.

We did not anticipate hypofrontality as we were interested in examining dlPFC activity during the flow experience and support for the hypofrontality hypothesis comes from observation of reduced mPFC activity during flow. While we could not evaluate synchronization of reward and cognitive control networks with our test setup, we anticipated greater signal coherence between frontal and temporoparietal regions during the experience of flow. Furthermore, we also anticipated moderate alpha activity in the flow condition compared to the go-signal task and the frustration conditions, as well as increased beta activity in frontal and temporoparietal regions. As we did not have midline frontocentral electrode coverage, we had no *a priori* hypotheses about theta activity in flow. A primary contribution of our study is a deeper understanding of candidate neural signatures obtained via lightweight wearable devices for real time monitoring of the flow state in closed-loop human–machine systems.

## II. Methods

### A. Participants

Ten adults (9 M, 1 F) provided informed consent and participated. All human participant procedures adhered to institutional guidelines set forth by Toyota Legal One at the institution leading the data collection. Following the experiment, participants were debriefed and provided detail on our research into relaxed concentration in mobility applications.

### B. Stop-Signal Task

Participants began the experiment on a laptop computer with a go-signal task (Fig. 1(a)). They were instructed to press ‘f’ when they saw a left-pointing arrow and ‘j’ when they saw a right-pointing arrow, as quickly as possible. Prior to each arrow, there was an inter-trial interval (ITI) drawn from an exponential distribution bounded by 500 and 2500 ms to avoid participants anticipating trial start times and locking into a rhythm. The arrow would display for a maximum of 650 ms while the task waited for a response. Immediately following a response, the task informed participants if they responded correctly or not with a red ‘X’ or a green check mark. Text would state that a participant was correct, pressed the wrong key, or did not press in time. Feedback would display for 1000 ms. The background was black while text and arrows were white; the text was presented in a sans-serif 72 pt font. There were 160 trials.

**Figure 1:**
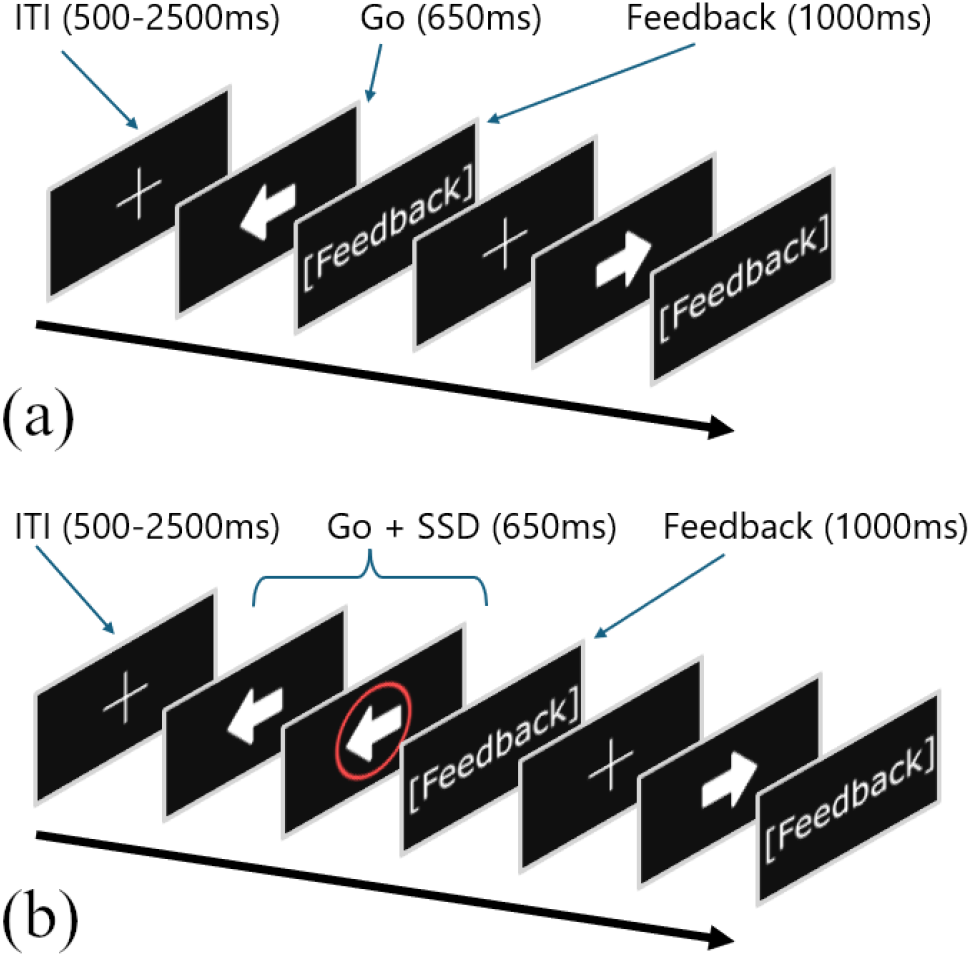
(a). Go-Signal Task. (b). Stop-signal task.

Following the go-signal task, participants engaged in the first of two stop-signal tasks (Fig. 1(b)); see [41], [42] for best practices. Similar to the go-signal task, participants had to press ‘f’ when they saw a left-pointing arrow and ‘j’ when they saw a right-pointing arrow. However, if a red circle appeared around the arrow, they were to withhold their response. Participants were informed if they responded correctly or not following each trial. For the trials with ‘stop’ signals, the task displayed an ‘X’ and informed the participant that they should not have pressed a key if they failed to withhold their motor response; the task displayed a green check mark with ‘Correct!’ as the accompanying text when participants successfully withheld a response. ITI and arrow presentation times were identical to the go-signal task. The stop-signal delay was initialized at 175 ms and increased or decreased on each ‘stop’ trial based on participant accuracy. For the ‘flow’ condition, a dynamic difficulty adjustment algorithm targeted 75% accuracy for ‘stop’ trials; for the ‘overload’ condition, the difficulty adjuster targeted 25% accuracy for ‘stop’ trials. There were 160 trials for each condition; a quarter of them included a ‘stop’ signal.

We did not counterbalance tasks as the go-signal task must precede the stop-signal tasks to build response automaticity. Furthermore, we wanted the flow condition to always follow the go-signal task so that the participant’s first experience of the stop-signal would be one of discovery. We also positioned the frustrating condition last as fatigue could contribute to a perception that it was overloading. Tasks were coded in Python 3.10.2 with PsychoPy version 2023.2.3, and administered on an MSI GS65 Stealth 9SF, with an Intel i9-9880H 2.3 GHz 8 core CPU, an NVIDIA Geforce RTX 2070 (laptop variant) GPU with base and boost clock speeds of 1215 MHz and 1440 MHz, and a 15.6” (1920x1080) 240 Hz display. We did not lock stimulus presentation to frame rate.

### C. Dynamic Difficulty Adjustment

The timing between ‘go’ and ‘stop’, the stop-signal delay (SSD), affects difficulty. As SSD increases, the probability of successfully inhibiting a response, *p*(*i*), decreases. Therefore, by adjusting SSD, we can make the participant feel in-control or frustrated. The difficulty adjuster targeted *p*(*i*) = 0.75 for flow and *p*(*i*) = 0.25 for frustration. Following a stop-signal trial, a *p*(*i*^*current*^) variable was updated based on the percentage of stop-signal trials the participant correctly responded to. We designed the difficulty adjustment algorithm so that the adjustment in the stop-signal delay was proportional to the difference between *p*(*i*^*target*^) and *p*(*i*^*current*^). The adjustment, *Adj*, in milliseconds, was drawn from an exponential function; see Equation (1). If *p*(*i*^*current*^) was exactly at the target following a stop-signal trial, no adjustment was made. Otherwise, the SSD adjustment between stop-signal trials could range between 16 ms and 67 ms, though in practice most adjustments were closer to 16 ms. We set constraints so that the earliest the stop-signal could appear would be 16 ms after the go signal, or four frames. Furthermore, the delay could not exceed the participant’s reaction time to the go stimuli determined during the go-signal task; the upper limit was set to 16 ms before the participant’s ‘go’ reaction time.

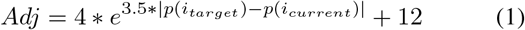

### D. Flow-State Scale - Short Form

Following each condition, participants filled out the short form of the Flow-State Scale (FSS) [43], [44], which queries nine proposed dimensions of flow [1], e.g., loss of sense of time, feeling in control of the situation, on a 7-point Likert scale; see Table I. No questions are reverse-coded.

**Table 1:** Short form of the FSS on a 7-point Likert scale.

| No. | Question |
| --- | --- |
| 1 | I feel just the right amount of challenge. |
| 2 | My thoughts/activities run fluidly and smoothly. |
| 3 | I do not notice time passing. |
| 4 | I have no difficulty concentrating. |
| 5 | My mind is completely clear. |
| 6 | I am totally absorbed in what I am doing. |
| 7 | The right thoughts/movements occur of their own accord. |
| 8 | I know what I have to do each step of the way. |
| 9 | I feel that I have everything under control. |
| 10 | I am completely lost in thought. |

### E. EEG Data Collection and Pre-processing

We used a lightweight commercial dry-contact EEG device, the Muse 2 [39], for cortical measurements. The Muse 2 collects EEG data at 256 Hz with 12-bit resolution with two frontal passive electrodes, AF7 and AF8, two temporoparietal passive electrodes, TP9 and TP10, one reference sensor at FpZ, and two bridged grounds at Fp1 and Fp2. It is also equipped with an accelerometer and gyroscope to monitor head translation and rotation. Smartphone communication with the Muse 2 occurs over Bluetooth^®^. The Muse^®^ app does not provide raw data, but a 3^rd^ party app, Mind Monitor [45], does. Prior to the task, electrodes, the participant’s forehead, and the area behind their ears were swabbed with alcohol pads (70%). The participant then put the headset on, wrapping it behind their ears and adjusting for a snug fit. The administrator instructed participants on how to position the headset to approximate good electrode placement based on the 10-20 system [46]. We evaluated electrode contact and signal quality before each condition and made adjustments as necessary.

Data were processed with custom scripts using Python 3.12.0, NumPy 1.26.3, SciPy 1.12.0, Pandas 2.2.0, and Scikit Learn 1.5.2. Signals were filtered with an IIR notch filter, 60 Hz with q = 30.0, to account for potential line noise. In addition, we noticed resonant spectral peaks near 22 Hz and 44 Hz in all channels but prominently in TP10. The source of these resonances is not attributable to aliasing from the device sample rate or Bluetooth^®^ frequency range. We also recreated these artifacts in multiple environments and could not attribute them to environmental noise; they are assumed to be an inherent characteristic of the device. Therefore, we also filtered signals with IIR notches at 22 Hz and 44 Hz with q = 10.0. Finally, we filtered signals with a Butterworth band pass filter with cutoffs at 1 and 50 Hz and an order of 4.

Next, translation and rotation parameters and their squared values were regressed from each electrode signal to remove variance due to head motion. Then, data were re-referenced to a common average reference to remove voltage bias from the frontal electrodes being closer to the FpZ reference than the temporal electrodes. Signal data were detrended and independent components analysis was performed with FastICA from Scikit Learn. Components that captured ocular noise were manually identified and regressed from all channels.

### F. Wavelet Analyses

The continuous wavelet transform (CWT) represents a signal in time and frequency. Conceptually, this can be achieved by convolving a signal with scaled and translated versions of a ‘mother’ wavelet function. These wavelets are localized in frequency and time. One way to think about wavelet convolution is as a spectral filter; the squared magnitude of the convolved signal provides information on power over time at the frequency targeted by a specific wavelet. Thus, CWT coefficients in plots indicate the strength of the signal at different frequencies and time positions; the CWT is particularly useful for analyzing EEG signals with dynamic characteristics where spectral power changes rapidly. Wavelet Transform Coherence (WTC) provides a time-frequency evaluation between two signals by measuring the local correlation between their wavelet transforms. It is useful for identifying regions with common power as well as identifying phase synchronization between non-stationary signals. For equations and discussion, see [47]–[49].

We extracted 1.25 second, 320-sample epochs for each trial; they included 500 ms before the ‘go’ signal and 750 ms after the ‘go’ signal. We manually evaluated them for artifacts and excluded an average of 6.67% of epochs per participant, *SD* = 4.70%. Epochs were detrended and normalized prior to wavelet transformation. We used a Morlet wavelet with six cycles as our mother wavelet. Rather than working with scale/octave divisions, we evaluated frequencies ranging from 1 Hz to 60 Hz in steps of 1 Hz. CWTs were calculated with Python 3.12.0 and PyCWT 0.4.0b [50]. To account for bias towards larger periods that can occur in wavelet analysis, we rectified the power spectrum [51]. We also calculated WCTs between AF7-AF8, AF7-TP9, AF8-TP10, and TP9-TP10.

### G. Statistical Analysis

The Wilcoxon Signed-Rank test [52] determines significant differences between paired observations when data is not normally distributed; as EEG data is far from a Gaussian process, we employed non-parametric tests. For each condition, for each participant, and for each electrode, we computed an average CWT matrix. These within-participant average matrices then served as input for the repeated-measures group analysis, which looked for differences between conditions. We also compared correctly inhibited to uninhibited motor responses on the stop-signal trials of the flow condition to evaluate spectral dynamics of performance lapses.

This was a limited exploratory analysis, so we evaluated differences at both the *p≤* 0.01 level and after false-discovery-rate correction [53] for multiple comparisons, *q* = 0.05. However, while significant at the *p≤* 0.01 level, no results survived FDR correction. Other tests were evaluated at *α* = 0.05. We performed calculations with Python 3.12.0, NumPy 1.26.3, SciPy 1.12.0, Pandas 2.2.0, and Statsmodels 0.14.4.

## III. Results

### A. Perception of Flow Across Conditions

We conducted a repeated measures ANOVA to examine differences across the experimental conditions in subjective report of the flow experience via the FSS. The overall main effect of condition was not statistically significant, *F*(2, 18) = 3.18, *p* = 0.06. However, post hoc comparisons using the Tukey Honestly Significant Difference test revealed a significant difference between the flow and frustration conditions (*t* = −2.39, *p* = 0.04), while the differences between the stop-signal conditions compared to the go-signal task did not reach significance (*t* = −0.39, *p* = 0.70; *t* = 1.63, *p* = 0.14). These findings suggest a specific contrast between flow and frustration despite the absence of a significant overall effect.

We initially hypothesized participants would report a greater sense of flow for the stop-signal task than the go-signal task, thinking that the go-signal task would be boring. However, we cannot label the go-signal task as a ‘Boredom’ condition in the absence of differences in subjective experience between tasks. Therefore, for the spectral analysis, we attributed differences in the contrasts between the go-signal condition and the stop-signal flow condition to task rather than psychoaffective state differences. Regardless, as the flow and frustration conditions were perceived differently, we attributed EEG differences between those conditions to psychoaffective state differences.

### B. Performance in Flow and Overload Conditions

Participant performance was tracked over successive trials for the flow and frustration conditions, during which task difficulty was continuously adjusted (Fig. 2). As intended, performance fluctuated around the targets set for the flow condition, and generally converged toward the target equilibrium as the task progressed. For the frustration condition, performance was variable and hovered near the intended equilibrium point. Three outliers performed better than expected on ‘stop’ trials. However, follow-up revealed that the performance of two of these outliers was lower on trials without a ‘stop’ signal (78% and 86% compared to the *∼*95% accuracy of other participants). Furthermore, their individual FSS scores were lower for this condition than the flow condition. Thus, these individuals likely adjusted their strategy to wait longer for the ‘stop’ signal on all trials, and they personally accepted failure on some of the ‘go’ trials to do better on the ‘stop’ trials. Overall, we interpreted performance results, alongside the subjective reports on the FSS, to validate that the flow condition elicited more of a flow experience than the frustration condition. The overall average SSD was 240 ms.

**Figure 2:**
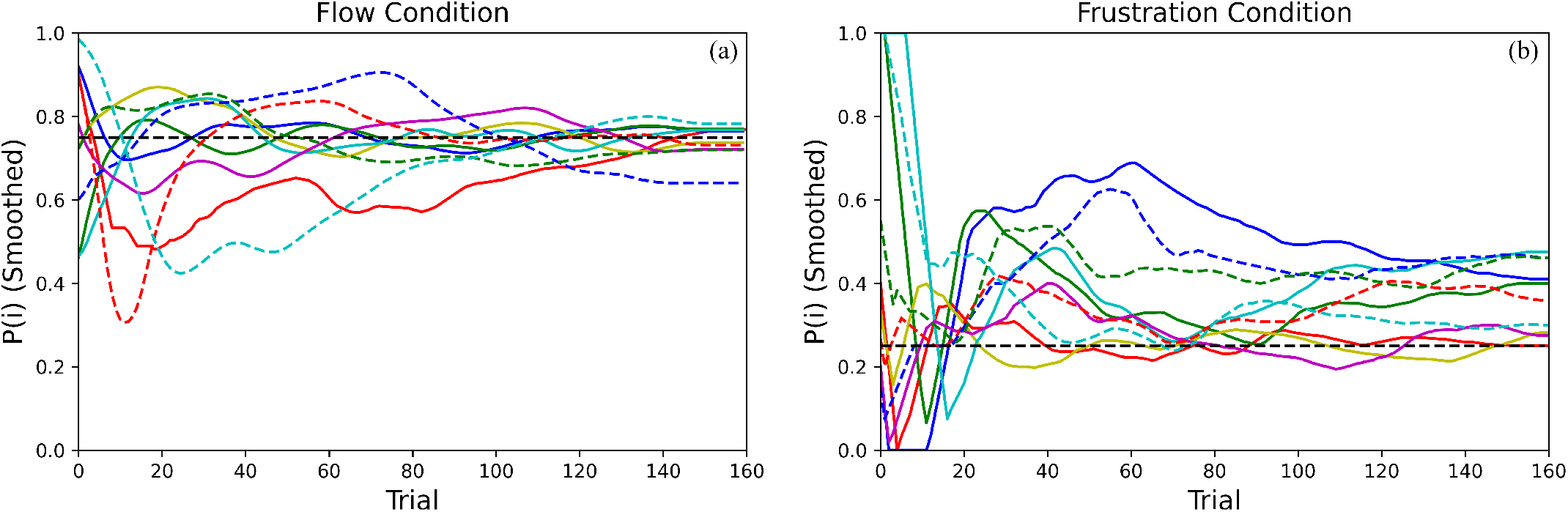
Tracking proportion of ‘stop’ trials where participants responded correctly. Horizontal dashed black lines indicate target probability of inhibition, *P* (*i*). (a) Individual traces tracking *P* (*i*) for flow. (b) Individual *P* (*i*) traces for frustration.

### C. Comparing the Go-Signal and Stop-Signal Tasks

Compared to the go-signal task, participants exhibited lower beta (18 − 30 Hz) coherence between AF7 and AF8 approximately 300 ms prior to the onset of the ‘go’ stimulus in the flow condition of the stop-signal task. There was also lower beta (14 − 18 Hz) coherence between AF7 and AF8 in the flow condition in-between the ‘go’ signal and the average SSD, as well as decreased frontal beta (18 − 20 Hz) coherence immediately after the average SSD (Fig. 3(a)-(d)).

**Figure 3:**
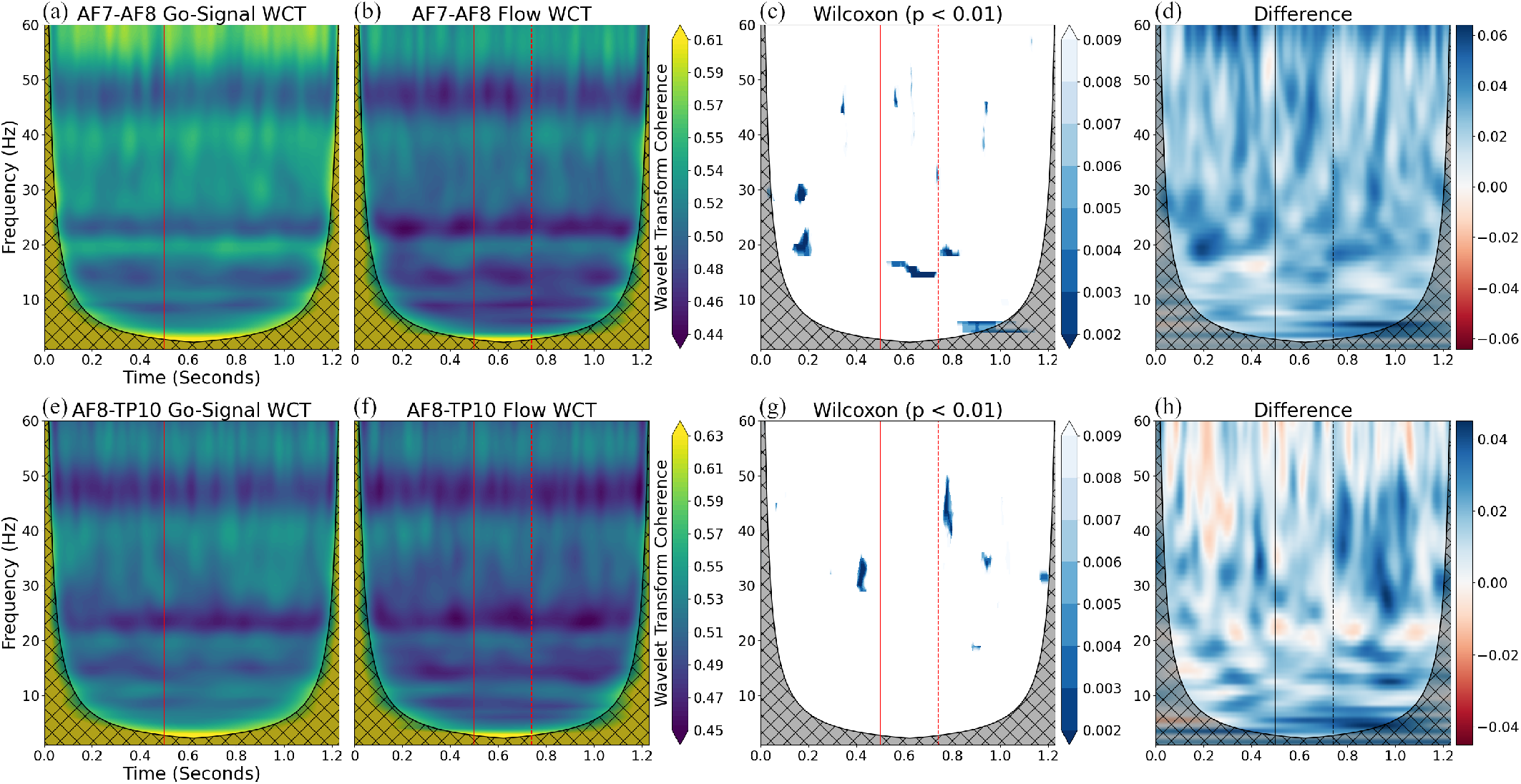
Spectral coherence for the go-signal task and the stop-signal flow condition. (a) Average AF7-AF8 go-signal WTC. (b) Average AF7-AF8 WTC for the flow condition. (c) Significant AF7-AF8 coherence differences between conditions, *p≤* 0.01. (d) AF7-AF8 WTC difference matrix (go-signal minus flow). (e) Average AF8-TP10 WTC for the go-signal task. (f) Average AF8-TP10 WTC for the flow condition. (g) Significant differences in AF8-TP10 coherence between conditions. (h) AF8-TP10 WTC differences matrix (go-signal minus flow). Solid lines indicate the ‘go’ signal; dashed lines indicate the average SSD.

In addition, in the stop-signal flow condition compared to the go-signal task, there was decreased gamma (30 − 35 Hz) coherence between AF8 and TP10 approximately 80 ms prior to the ‘go’ signal and decreased gamma (38 − 50 Hz) coherence roughly 40 ms after the average SSD (Fig. 3(e)-(h)). There were also bouts of gamma decoherence between 30 and 40 Hz in the flow condition roughly 100 ms and 300 ms after the average SSD (Fig. 3(e)-(h)). As some of these coherence differences surround the mean timing of the ‘stop’ signal, and participants rated these conditions similarly via the FSS, the differences likely reflect task differences rather than neural signatures of flow.

### D. Comparing Correct and Incorrect Flow Task Trials

There was a reduction in TP10 gamma power (45 − 55 Hz) approximately 250 ms before the ‘go’ signal and 50 ms before the average SSD in ‘stop’ trials where participants failed to withhold their response during the flow condition (Fig. 4(a)-(d)). Furthermore, during flow, participants exhibited decreased AF7 beta (23 − 30 Hz) power 250 ms before the ‘go’ stimulus and decreased gamma (38 − 42 Hz) power 100 ms before the ‘go’ stimulus for trials where they responded erroneously to the ‘stop’ compared to trials where they withheld responses (Fig. 4(e)-(h)). We also observed decreased AF7 gamma (50 − 60 Hz) roughly 50 ms before the average ‘stop’ stimulus timing immediately followed by elevated gamma (45 − 50 Hz) roughly 25 ms before the average ‘stop’ signal for the ‘stop’ trials where participants made an error. Elevated beta (18 − 20 Hz) roughly 100 ms after the average ‘stop’ stimulus timing was also observed for trials where participants failed to withhold their motor response (Fig. 4(e)-(h)). There was also decreased gamma (49 − 54 Hz) coherence between electrodes AF7 and TP9 around the presentation of the ‘stop’ signal during the flow condition when comparing ‘stop’ trials that participants incorrectly and correctly responded to (Fig. 4(i)-(l)).

**Figure 4:**
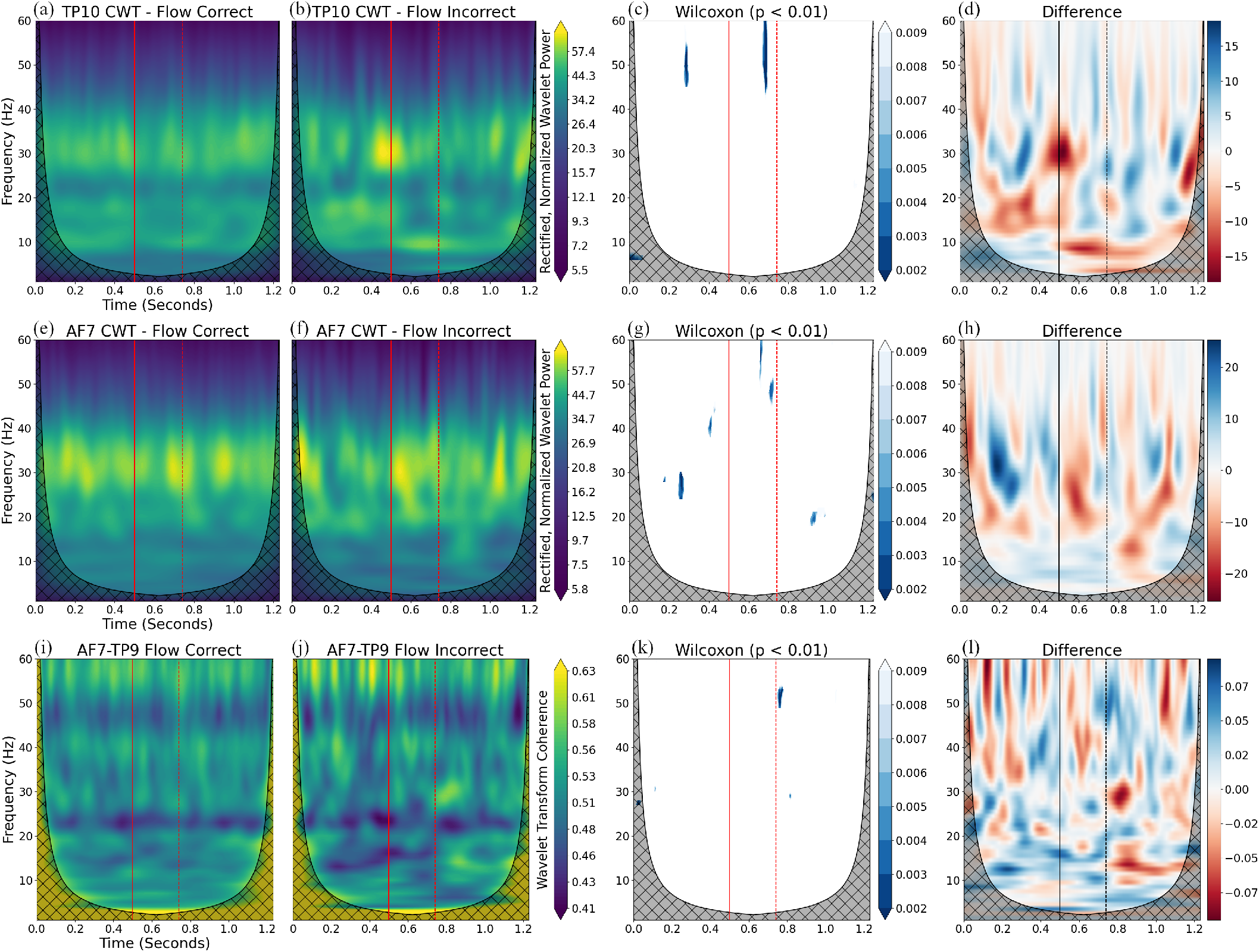
Spectral dynamics during correct and incorrect responses to ‘stop’ trials during the flow condition. (a) Average TP10 CWT for trials when participants responded correctly. (b) Average TP10 CWT for when participants responded incorrectly. (c) Significant TP10 CWT differences between success and failure at *p≤* .01. (d) Differences between the average TP10 CWT arrays (correct minus incorrect). (e) Average AF7 CWT power during ‘stop’ trials when participants responded correctly. (f) Average AF7 CWT power for ‘stop’ trials where participants failed to withhold a response. (g) Significant differences. (h) AF7 CWT difference matrix (correct minus incorrect). (i) Average AF7-TP9 WTC for flow condition trials where participants responded correctly to the ‘stop’ signal. (j) Average AF7-TP9 flow condition trials where participants responded incorrectly to the ‘stop’ signal. (k) Significant differences. (l) Differences between the average AF7-TP9 WTC arrays (correct minus incorrect). Solid lines indicate the ‘go’ signal; dashed lines indicate the average SSD.

### E. Spectral Dynamics in Flow and Frustration

Compared to the flow condition, there was greater beta (14 − 20 Hz) power between the ‘go’ and ‘stop’ signals in the frustration condition at TP9 and TP10 (Fig. 5). The peaks of these clusters were approximately 50 ms before the average timing of the ‘stop’ signal. There was also greater temporal beta power in the frustration condition roughly 350 ms after ‘stop’ stimulus presentation.

**Figure 5:**
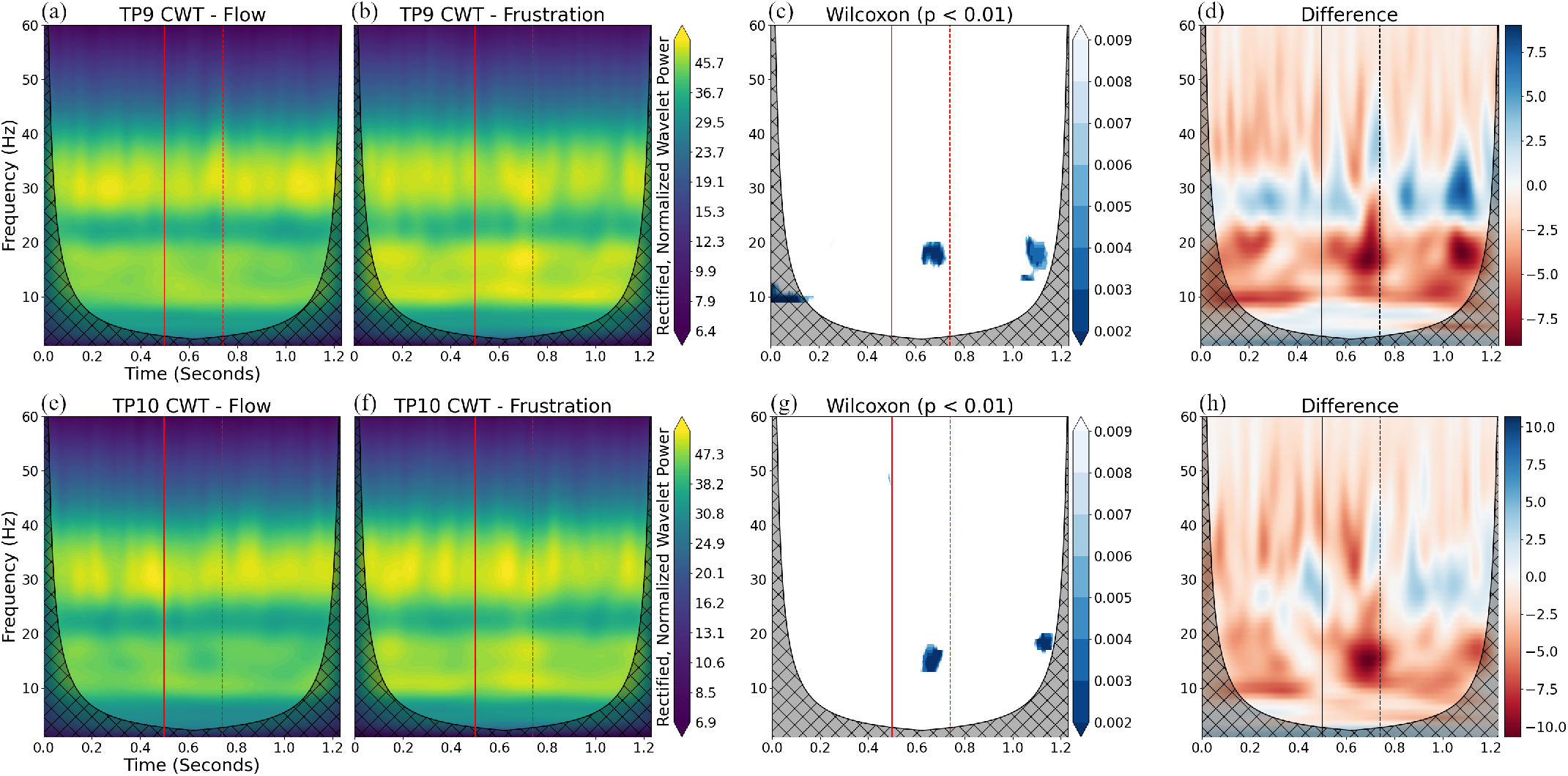
TP9 and TP10 spectral dynamics. (a) Average TP9 CWT during flow trials. (b) Average TP9 CWT during frustration trials. (c) Significant differences between conditions at *p≤* .01. (d) TP9 difference matrix (flow minus frustration). (e) Average TP10 CWT during flow trials. (f) Average TP10 CWT during frustration trials. (g) Significant differences. (h) TP10 difference

## IV. Discussion

Here, we investigated spectral dynamics that accompany a flow experience during a stop-signal task, contrasting them with dynamics observed during go-signal and frustrating overload conditions. As predicted, subjective ratings on the short form FSS diverged between the flow and overload blocks, suggesting that the difficulty adjustment separated a seamless, intrinsically rewarding state from an aversive, high cognitive load state. However, the go-signal task was not perceived as ‘boring’ compared to the flow condition as we had hoped; therefore, we were only able to contrast flow with frustration and not flow with boredom. Guided by recent work on the neural signatures of flow, we anticipated that the experience of flow would be accompanied by strong coherence between frontal (AF7/AF8) and temporoparietal (TP9/TP10) areas, signaling functional coupling of central executive regions. In addition, we anticipated moderate alpha power and elevated beta power across electrodes during flow relative to the other conditions, reflecting relaxed but moderate effort.

### A. Spectral Dynamics Related to Task Differences

In comparing the stop-signal task under the flow condition to the go-signal task, we found notable differences in interregional EEG coherence. In the go-signal task, participants exhibited lower beta coherence between left and right frontal electrodes prior to the ‘go’ signal and in the window between the ‘go’ signal and the SSD. Similarly, gamma coherence between right frontal and temporoparietal electrodes was reduced in the flow condition immediately around the ‘go’ signal and shortly after the average SSD. In general, participants exhibited less synchronized activity between frontal regions and between frontal and temporoparietal regions in the stop-signal task compared to the go-signal task.

We were unable to speculate on EEG differences between flow and boredom with the present data; the coherence reductions are likely better attributed to the required inhibitory control and timing uncertainty added into the stop-signal task. Prior EEG study of response inhibition has observed somewhat of an opposite pattern during No-Go trials of a Go/No-Go task, with interhemispheric frontal coherence (F3–F4) increasing during successful response suppression [54]. Specifically, they found significantly higher prefrontal coherence in narrow frequency bands around 3.9 and 7.8 Hz approximately 250 − 450 ms after stimulus presentation; indicating that coordinated theta and alpha activity facilitate decision making and suppression of a motor response.

Perhaps, decreased beta coherence during the stop-signal task could reflect differences in timing between No-Go and Stop-Signal tasks. The ‘stop’ signal in No-Go trials is presented immediately at the start of the trial, whereas the ‘stop’ signal is delayed for stop-signal paradigms. The delay may require more flexible updating or switching between maintained rulesets. High beta in the dlPFC reflects active top-down control of the current cognitive state; and frontal beta synchrony is associated with rulesets [55]. Weakening prefrontal beta coupling could allow for new information to override maintained information. Decreased coherence could reflect increasing executive functioning demands, and switching from a top-down predictive coding state to a bottom-up state where there is uncertainty about the action to take [56]– [59]. Regardless, interpretation remains unclear and requires further investigation.

### B. Spectral Dynamics of Flow and Overload

Frustration was marked by elevated temporoparietal beta power (14-20 Hz) between the onset of the ‘go’ signal and the expected timing of the ‘stop’ signal, whereas flow showed a tempered beta profile. The function of TPJ beta is not well-established compared to PFC, but elevated beta in TPJ could indicate engagement in cognitive or perceptual integration processes. The TPJ sits at a crossroads of sensory inputs and combines sensory processing with attentional orienting [60]. High beta suggests that the TPJ possibly maintains an expectation. In our frustrating scenario, TPJ beta could reflect the strain of tracking a salient, infrequent signal. Elevation in beta power may reflect oscillatory activity linked to cognitive effort and set maintenance [61]–[63]. Furthermore, beta oscillations are not idle rhythms, but exhibit transient bursts that carry topdown information relevant for discrete cognitive operations [64]. Compared to frustration, the flow block showed tempered beta activity, consistent with the idea appropriate challenge elicits efficient but not excessive engagement of executive circuitry, aligning with the Neural Proficiency perspective.

We expected differences in alpha dynamics between conditions as alpha oscillations relate to attention; alpha power tends to decrease in task-relevant regions as cognitive load increases [34], [35]. However, we did not find alpha differences between our conditions. Moderate alpha activity in frontal regions has previously been linked to the experience of flow [7], [37]. However, we were unable to corroborate these results. It could be that increasing cognitive workload through arithmetic has a different effect on alpha activity than managing a stop-signal delay. Furthermore, participants experienced a similar level of flow for the go-signal task as they did for the flow condition of the stop-signal task. Thus perception of difficulty may not have been substantially different between conditions to observe a moderate alpha profile in flow.

### C. Spectral Dynamics of Efficient Task Performance

Our analyses revealed spectral patterns that distinguished successful from failed inhibitory responses. Brief lapses in beta and gamma power foreshadowed errors. Specifically, on stop-signal trials where participants failed to withhold a response, we observed a reduction in beta power over the left dlPFC during the ITI before the ‘go’ cue. Decreased beta power in dlPFC during the ITI could indicate failure to “clear-out” information in working memory between trials for the trials where participants responded incorrectly. The drop in beta power could also reflect a lapse in attentional maintenance.

On error trials, we found a dip in left dlPFC gamma power before the expected stop signal. Error-related drops in dlPFC gamma power were followed by a late gamma burst before the average timing of the stop signal. Lower gamma power immediately followed by elevated gamma for the incorrect trials could also indicate a delay or timing mismatch in the bottom-up processing of stop-signal information entering into dlPFC. Indeed, prior literature suggests that timing of gamma burst activity can relate to successful task performance [62]. For our analysis, the late dlPFC gamma burst on error trials could reflect an abort signal that was dispatched too slowly.

An additional observation was that brief reductions in temporoparietal gamma power and left fronto-temporoparietal gamma coherence preceded failures to inhibit action. Reductions in gamma power and coherence suggest that the cortical assemblies governing executive control and attention momentarily disengage before erroneous responses. Transient desynchronizations and drops in spectral power align with established views of gamma bursts as signatures of working memory updating and prediction error [61], [62]. The TPJ detects salient events and the PFC suppresses or alters action plans; coupling between these regions in the gamma band may represent the communication of ‘stop’ information and the mobilization of inhibitory control. Finally, transient spectral events in the beta band also followed the outcomes of stop trials. We found that trials with failed inhibition elicited a spike of beta power about 100 ms after the stop-signal time in the AF7 electrode that was higher than in successful trials. One interpretation is that this late beta burst represents a post-error “clear-up” or compensatory process [56].

Practically, millisecond-scale markers are candidate features for closed-loop attention monitoring systems, e.g., determining if a driver will respond to dynamic road conditions. Elevated TPJ gamma could reflect active processing of salient information within attention networks; gamma could be a sign of local neural assembly firing [65], [66] and gamma bursts may indicate salient sensory input processing and attentional shifts [67]. During demanding tasks, if an unexpected event occurs, TPJ gamma may spike, indicating that attentional regions are rapidly processing the event. Therefore, elevated gamma activity observed in ‘stop’ trials where participants responded matrix (flow minus frustration). Solid lines indicate the ‘go’ signal; dashed lines indicate the average SSD. correctly may reflect a neural signature of timely, accurate processing of environment changes. This pattern of brain activity could be a viable feature to automatically determine if a human will respond to a change in their environment in time, e.g., a driver responding to a railroad crossing signal.

### D. Limitations

Several constraints temper generalizability. The sample was small and demographically narrow (90% male engineers), limiting power and external validity. In addition, only four dry-contact EEG electrodes were available, restricting spatial resolution and preventing assessment of deep reward structures central to the Synchronization Hypothesis. Task order was also fixed, so fatigue or expectation could confound condition differences. Finally, device artifacts at 22 Hz and 44 Hz required aggressive filtering, impacting genuine neural signal.

## V. Conclusion

Balanced challenge produces a distinct neurophysiological signature of flow marked by low beta power, whereas mild frustration is marked by beta power elevations indicative of heightened cognitive effort. Pre-stimulus collapses in right temporoparietal gamma power and frontal-temporoparietal gamma coherence precede task errors in flow, highlighting candidate features for real time vigilance monitoring in closed loop human–machine systems. While limitations temper generalizability, the results indicate that millisecond-scale beta and gamma patterns capture elements of flow and frustration.

## Acknowledgment

The authors would like to thank Jae Seung Lee, Yukiko Ohnishi, and Yasushi Dohnoue for their discussions and support of this project.

